# Employing biological sex as a primary variable can restrict our understanding of behavioral diversity (and sex differences) in psychostimulant activity

**DOI:** 10.1101/2024.11.06.622330

**Authors:** Gabriella Yao, Martin O Job

## Abstract

**Background:** In addition to representing distinct biological sex groups, males and females are also individuals expressing behavioral diversity. Our MISSING (Mapping Intrinsic Sex Similarities as an Integral quality of Normalized Groups) model suggests that, with respect to behavior, grouping according to the individual reveals groups with differences that exceed biological sex-related differences, but this needs further clarification. We hypothesized that, compared to the current model (grouping by biological sex), the MISSING model (grouping by individual attributes) was the more effective grouping strategy to identify behavioral diversity.

**Methods:** We conducted experiments in rats to determine the locomotor activity (in 90 min) following intraperitoneal injections of saline (n = 12 males, n = 11 females) and cocaine (n = 8 males, n = 11 females). For the current model, we compared males versus females using unpaired t-tests. For the MISSING model, we identified clusters of individuals (males and females) using normal mixtures clustering analysis of several behavioral variables and employed unpaired t-tests to compare clusters and Two-way ANOVA to determine if there were any SEX by cluster interactions. For both models, we employed linear regression analysis to compare relationships between variables and Two-way repeated measures ANOVA to analyze locomotor activity time course.

**Results:** For both the saline and cocaine groups, the MISSING model identified two behavioral clusters with differences that exceeded any differences due to biological sex.

**Conclusions:** The MISSING model suggests that employing biological sex as a primary variable can obscure our understanding of sex and individual differences in psychostimulant activity.

## Introduction

The importance of understanding sex differences in the effects of drugs, including addictive substances, cannot be overemphasized (Becker, 1999; Becker et al., 2016; Becker and Chartoff, 2019; Becker and Hu, 2008; Becker and Koob, 2016; Beery and Zucker, 2011; Miller et al., 2017; Prendergast et al., 2014; Shansky and Woolley, 2016; Zucker and Beery, 2019). However, males and females are also individuals. Individuals, regardless of biological sex, express behavioral diversity. Thus, males and females also have behavioral group identities independent of their biological sex. However, a determination of which of these identities (biological sex versus behavioral group) would be a more effective grouping strategy for the understanding of individual and sex differences has not been done.

To account for both individual and biological sex differences, we developed a model which we termed the MISSING (Mapping Intrinsic Sex Similarities as an Integral quality of Normalized Groups) model (see Figure 1B), compare this to the current model (Figure 1A). The MISSING model which we validated in previous reports (Job, 2024; Showell and Job, 2024; Tigano and Job, 2024), proposes that 1) there are no sex differences when we compare males and females within the same behavioral group, 2) sex differences occur when we compare males and females from different groups and is tantamount to a group effect and not a biological sex effect *per se*, and 3) even if/when we detect sex differences between males and females in the same behavioral group, these differences will not be as significant as differences between males and females from different groups. The MISSING model suggests that grouping individuals (independent of biological sex) reveals groups with distinctions exceeding any differences we can realize if we group by biological sex, but this needs to be further clarified.

**Figure 1:**
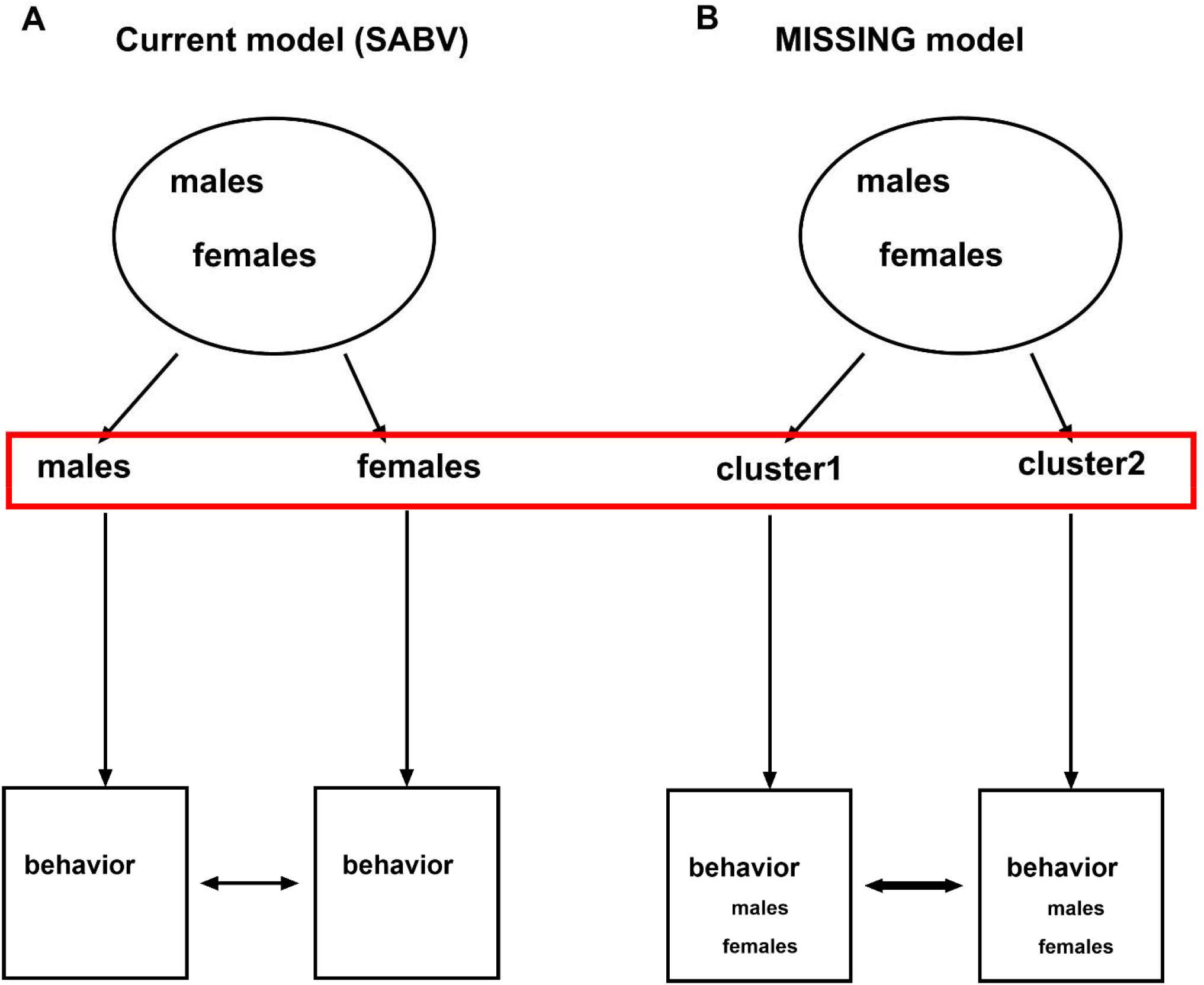
The current model and the MISSING model: model descriptions. Fig A represents the current model (with the primary variable as SABV – sex as a biological variable). From a pool of both sexes, males and females are separated into groups and their behaviors are compared to determine if there are sex differences. Fig B is the new Mapping Intrinsic Sex Similarities as an Integral quality of Normalized Groups (MISSING) model. For MISSING, biological sex is not a primary variable, behavioral group is the primary variable. From a pool of both sexes, males and females are NOT separated into groups, rather the whole pool of individuals undergoes normal mixtures clustering analysis of several variables to determine which groups/clusters, irrespective of biological sex, the individual belongs. It is only afterwards that males and females within a cluster are compared. This is proposed to be a more accurate assessment of sex differences (males versus females within a cluster and not males versus females generally). Comparisons of males and females that do not belong to the same clusters will overestimate sex differences because those sexes already belong to distinct groups and the sex difference, in this case, will be tantamount to the group difference. In this study we wanted to determine which model (Fig A or Fig B) would reveal groups that were more behaviorally distinct (males versus females OR cluster versus cluster).

Thus, we hypothesized that, compared to the current model (grouping by biological sex), the MISSING model (grouping by individual attributes) was the more effective grouping strategy to identify behavioral diversity. Behavioral diversity is an important concept to understand if we are to explain individual differences in the effects of addictive substances. We proceeded to test this hypothesis by conducting experiments to determine the locomotor activity of male and female Sprague Dawley rats following vehicle and cocaine injection at a dose = 15 mg/kg via the intraperitoneal route. We included vehicle controls and this cocaine dose because the MISSING model has not been validated for these conditions. We obtained locomotor activity variables similar to previous studies (Job, 2024; Tigano and Job, 2024) and we compared these variables (and the relationships between these variables) to determine which grouping strategy (males versus females OR cluster versus cluster) revealed groups that were truly distinct behaviorally. Our methods, results and discussions are below.

## Methods

Experiments and animal care were in accordance with the Institute of Animal Care and Use Committee of Emory University and followed the guidelines outlined in the National Institutes of Health (NIH) *Guide for the Care and Use of Laboratory Animals*. We employed a total of forty-two (42) adult male and female adult Sprague-Dawley rats that weighed between 225 and 275g at the time of purchase from Charles River Laboratories (Wilmington, MA). They were acclimatized to the housing facility for at least one week before any experimental procedures were conducted. They were provided with rat chow and water *ad libitum* and maintained on a 12-hour light: dark cycle (lights on at 7 am). We conducted all experiments between 10:00 AM and 6:00 PM.

### Experimental groups

see Table.

Experimental design: There were two groups – saline group and cocaine group. There were n = 12 males and n = 11 females in the saline group and there were n = 8 males and n = 11 females in the cocaine group. After 30 min of baseline activity measurements, the saline group was injected with cocaine 0 mg/kg via the intraperitoneal route while the cocaine group was injected with cocaine 15 mg/kg via the same route and total distance traveled in cm over 90 min (after injections) were assessed.

|  | Saline group | Cocaine group |
| --- | --- | --- |
| males | N = 12 | N = 8 |
| females | N = 11 | N = 11 |
| total | N = 23 | N = 19 |

### Drugs

Cocaine (0 mg/kg, saline, vehicle control) and 15 mg/kg via the intraperitoneal route.

### Assessments of locomotor activity

These were done as in previous studies (Job, 2024, 2016; Job et al., 2014, 2013, 2012; Job and Kuhar, 2017, 2012; Kuhar and Job, 2017; Tigano and Job, 2024).

### Variables

There were three variables assessed – baseline activity, saline/cocaine activity and saline/cocaine activity normalized-to-baseline activity (NBA). Baseline activity was estimated as distance traveled (cm) in 30 min just prior to saline and/or cocaine injection. Saline and/or cocaine activity was assessed as distance traveled in 90 min after saline and/or cocaine injection. Saline activity NBA and cocaine activity NBA were calculated as shown in the equations below:

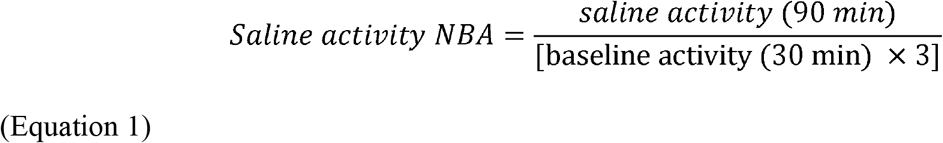

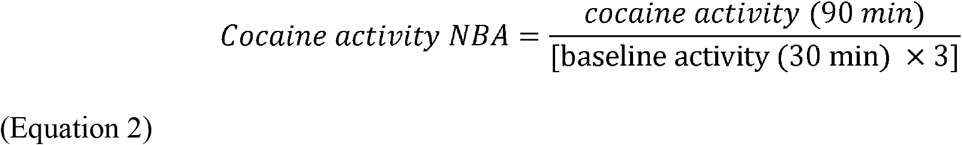

Because saline/cocaine activity occurred over 90 min and baseline was only 30 min, normalization involved multiplying baseline activity by 3. For more details regarding these variables, see (Job, 2024; Tigano and Job, 2024), see also (Showell and Job, 2024).

### Statistical analysis

We employed GraphPad Prism v 10 (GraphPad Software, San Diego, CA), SigmaPlot 14.5 (Systat Software Inc., San Jose, CA) and JMP Pro v 17 (SAS Institute Inc., Cary, NC) for statistical analysis. Grubb’s test was used to determine if there were any significant outliers. Data were expressed as mean ± SEM. For the current model, we separated groups by biological sex (males versus females) and employed unpaired t-tests to determine if there were differences in the average of variables. We employed linear regression to determine if there were differences between biological sex groups with regards to the relationships between variables. For analysis of time course data, we employed Two-way repeated measures ANOVA with factors as SEX (males and females) and time to determine if there was a SEX × time interaction, a main effect of SEX and a main effect of time. For the MISSING model, we employed normal clustering of several variables (baseline activity, saline/cocaine activity and saline/cocaine activity NBA) for every individual, regardless of sex, to identify distinct clusters. As with the current model, we employed unpaired t-tests to determine if there were differences between behavioral clusters in the average of variables. We employed linear regression to determine if there were differences between clusters with regards to the relationships between variables. For analysis of time course data, we employed Two-way repeated measures ANOVA with factors as cluster and time to determine if there was a cluster × time interaction, a main effect of cluster and a main effect of time. To determine if there was sex differences in the time course data within the same cluster and between different clusters, we employed a Two-way ANOVA to determine if there was a SEX × time interaction. Statistical significance was set at p < 0.05 for all analyses with Tukey’s post hoc test employed when significance was detected.

## Results

### Identification of biological sex groups

In line with the current model, subjects were grouped as males and females based on external genitalia. For the saline group, we had n = 12 males and n = 11 females. For the cocaine group, we had n = 8 males and n = 11 females.

### Identification of clusters (MISSING model)

For the saline group, normal mixtures clustering analysis of baseline activity, saline activity and saline activity NBA yielded two clusters which we termed cluster1 and cluster2. Cluster1 consisted of n = 7 males and n = 5 females whereas cluster2 consisted of n = 5 males and n = 6 females (Figure 2A-B).

**Figure 2:**
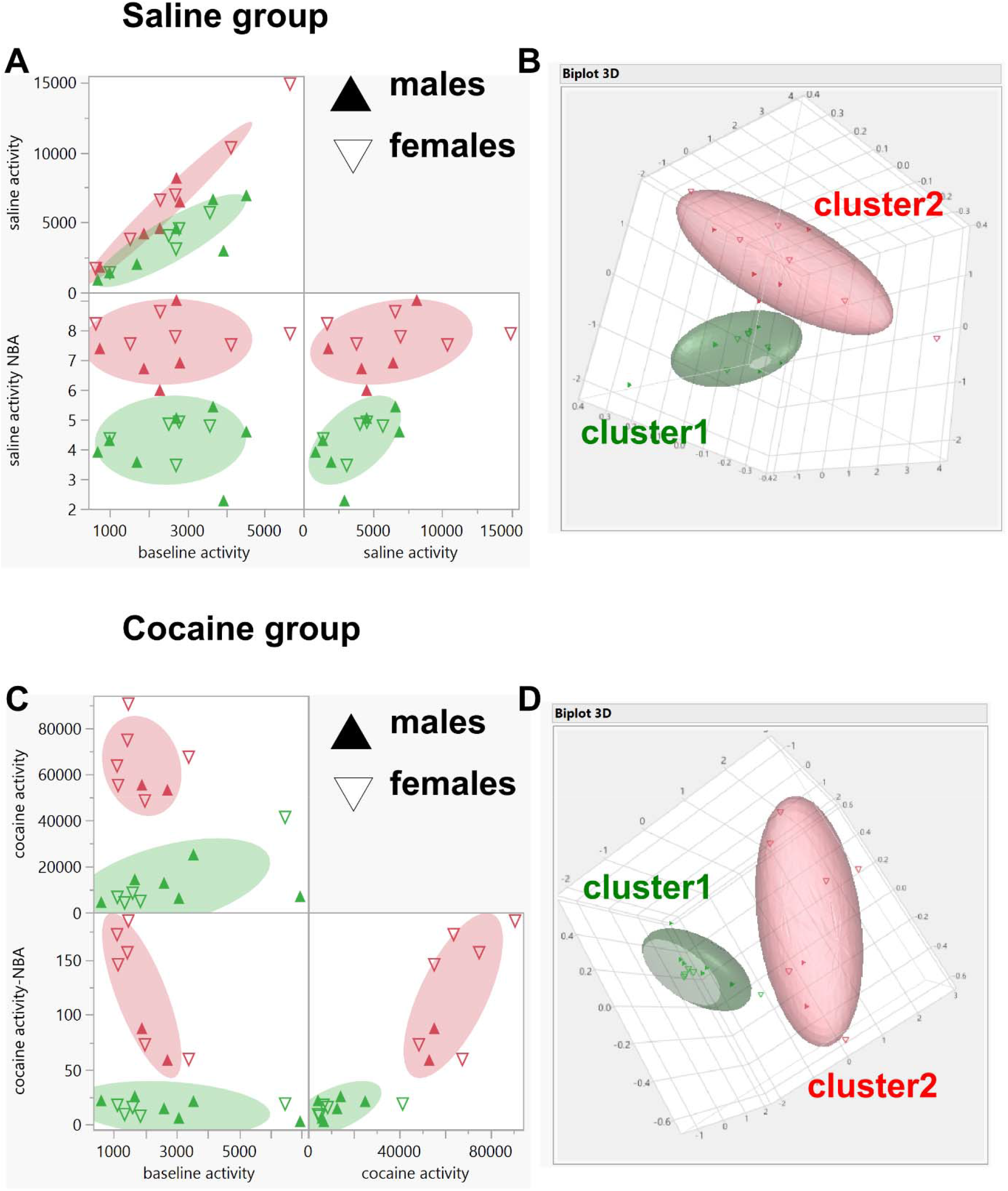
Identification of clusters (MISSING model) for the saline and cocaine groups. We conducted clustering analysis (normal mixtures) of baseline activity, saline activity and saline activity NBA for all individuals in the saline group and identified two clusters (Fig A-B). We conducted normal mixtures clustering analysis of baseline activity, cocaine activity and cocaine activity NBA for all individuals in the cocaine group and identified two clusters (Fig C-D). The closed triangles represent males while the open triangles represent females. The green and red colored data points represent individuals in cluster1 and cluster2, respectively.

For the cocaine group, normal mixtures clustering analysis of baseline activity, cocaine activity and cocaine activity NBA yielded two clusters which we termed cluster1 and cluster2. Cluster1 consisted of n = 6 males and n = 5 females whereas cluster2 consisted of n = 2 males and n = 6 females (Figure 2C-D).

### Model comparisons: saline group

#### Males versus Females

For the saline group, the baseline activity of males (n = 12) and females (n = 11) were 2382 ± 363 and 2681 ± 430 cm, respectively. The saline activity of males (n = 12) and females (n = 11) were 4208 ± 703 and 5759 ± 1195 cm, respectively. The saline activity NBA of males (n = 12) and females (n = 11) were 5.42 ± 0.54 and 6.37 ± 0.57, respectively. Unpaired t-tests revealed no significant differences between males and females for baseline activity (P = 0.5997, Figure 3A), saline activity (P = 0.2661, Figure 3B) and saline activity NBA (P = 0.2388, Figure 3C). For the relationship between baseline activity and saline activity, linear regression analysis revealed that there were no differences between males and females for slope (F 1, 19 = 3.868, P = 0.0640) and y-axis intercept (F 1, 20 = 1.471, P = 0.2394), see Figure 3D. For the relationship between baseline activity and saline activity NBA, linear regression analysis revealed that there were no differences between males and females for slope (F 1, 19 = 0.2299, P = 0.6371) and y-axis intercept (F 1, 20 = 1.413, P = 0.2484), see Figure 3E. For the relationship between saline activity and saline activity NBA, linear regression analysis revealed that there were no differences between males and females for slope (F 1, 19 = 0.6048, P = 0.4463) and y-axis intercept (F 1, 20 = 0.5210, P = 0.4788), see Figure 3F.

**Figure 3:**
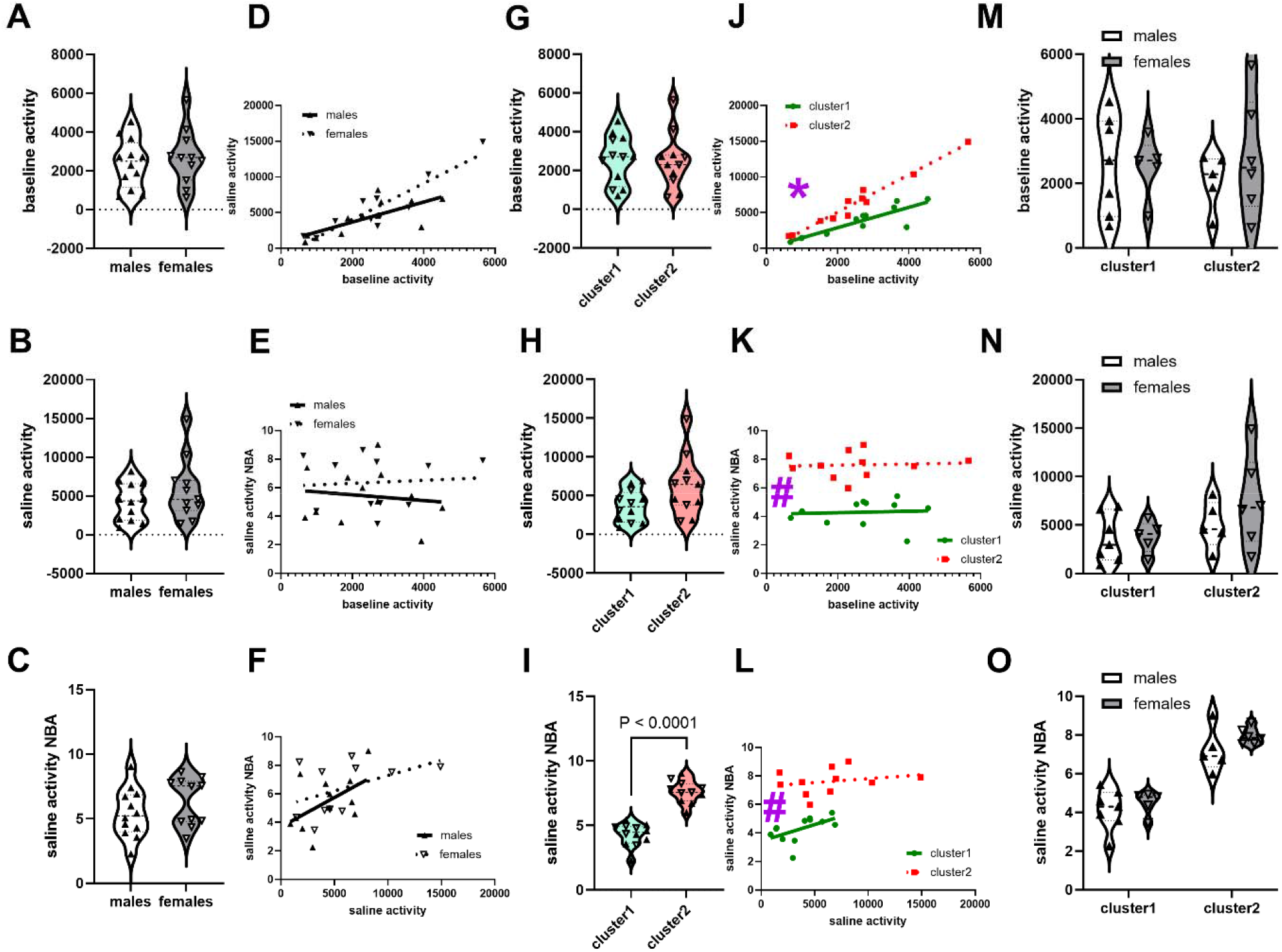
The behavioral diversity between clusters exceeds biological sex differences: saline group. Fig A-C show comparisons between males and females for baseline activity, saline activity and saline activity NBA, respectively, in accordance with the current model. Fig D-F, respectively, show the relationships between baseline activity and saline activity, baseline activity and saline activity NBA, and saline activity and saline activity NBA, for males and females, respectively. Fig G-I show comparisons between MISSING model-identified cluster1 and cluster2 for the same variables as in Fig A-C. Fig J-L, respectively, show the relationships between baseline activity/ saline activity, baseline activity/ saline activity NBA, and saline activity/ saline activity NBA, for cluster1 and cluster2. Fig M-O show comparisons of sex within and between clusters to determine if there was a SEX by cluster interaction (MISSING model). There were no differences between males and females for any variables/ variable relationships (Fig A-F). There were differences in variables/ variable relationships between clusters (Fig I-L), except for the baseline activity and saline activity variable (Fig G-H). Two-way ANOVA revealed no SEX by cluster interaction for all variables and no main effect of SEX for all variables (Fig M-O), but a main effect of cluster for saline activity NBA (P < 0.0001). In summary, there were more significant differences for variables/ variable relationships between clusters than between biological sex. The * and # show significant differences (P < 0.05) for slope and y-axis intercepts, respectively.

#### Cluster1 versus cluster2

The baseline activity of cluster1 (n = 7 males, n = 5 females) and cluster2 (n = 5 males, n = 6 females) were 2563 ± 363 and 2484 ± 436 cm, respectively. The saline activity of cluster1 and cluster2 were 3689 ± 595 and 6325 ± 1166 cm, respectively. The saline activity NBA of cluster1 and cluster2 were 4.29 ± 0.25 and 7.61 ± 0.26, respectively. Unpaired t-tests revealed no significant differences between cluster1 and cluster2 for baseline activity (P = 0.8886, Figure 3G) and saline activity (P = 0.0513, almost significant, Figure 3H) but revealed significant differences for saline activity NBA (P < 0.0001, Figure 3I). For the relationship between baseline activity and saline activity, linear regression analysis revealed that there were significant differences between cluster1 and cluster2 for slope (F 1, 19 = 16.37, P = 0.0007, Figure 3J). For the relationship between baseline activity and saline activity NBA, linear regression analysis revealed no significant differences (cluster1 versus cluster2) between slopes (F 1, 19 = 0.001101, P = 0.9739), but significant differences between y-axis intercepts (F 1, 20 = 79.62, P < 0.0001, Figure 3K). For the relationship between saline activity and saline activity NBA, linear regression analysis revealed that there were no differences between clusters for slope (F 1, 19 = 1.632, P = 0.2169), but there were significant differences between clusters with regards to the y-axis intercept (F 1, 20 = 63.28, P < 0.0001, Figure 3L).

We wanted to know if there were sex differences when we account for the clusters. For this, we employed a Two-way ANOVA with factors SEX (males and females) and cluster (cluster1, cluster2). For baseline activity (Figure 3M), we detected no SEX × cluster interaction (F 1, 19 = 0.4894, P = 0.4927), no main effect of cluster (F 1, 19 = 0.03309, P = 0.8576) and no main effect of SEX (F 1, 19 = 0.3128, P = 0.5825). For saline activity (Figure 3N), we detected no SEX × cluster interaction (F 1, 19 = 0.7203, P = 0.4066), no main effect of cluster (F 1, 19 = 3.748, P = 0.0679) and no main effect of SEX (F 1, 19 = 0.9547, P = 0.3408). For saline activity NBA (Figure 3O), we detected no SEX × cluster interaction (F 1, 19 = 0.3291, P = 0.5729), and no main effect of SEX (F 1, 19 = 2.196, P = 0.1548), but a significant effect of cluster (F 1, 19 = 80.34, P < 0.0001).

### Model comparisons: cocaine group

#### Males versus Females

The baseline activity of males (n = 8) and females (n = 11) were 2884 ± 666 and 2084 ± 481 cm, respectively, and unpaired t-tests revealed no difference between males and females with regards to this variable (P = 0.3310, Figure 4A). The cocaine activity of males (n = 8) and females (n = 11) were 22193 ± 7335 and 42417 ± 9501 cm, respectively. Comparisons of average cocaine activity using unpaired t-tests revealed no biological sex differences (P = 0.1331, Figure 4B). The cocaine activity NBA of males (n = 8) and females (n = 11) were 29.75 ± 10.19 and 78.58 ± 21.76, respectively. When we compared males and females using unpaired t-tests, we determined that there were no differences for this variable (P = 0.0895, Figure 4C). For the relationship between baseline activity and cocaine activity, linear regression analysis revealed that there were no differences between males and females for slope (F 1, 15 = 0.1506, P = 0.7034) and y-axis intercept (F 1, 16 = 2.155, P = 0.1615), see Figure 4D. For the relationship between baseline activity and cocaine activity NBA, linear regression analysis revealed that there were no differences between males and females for slope (F 1, 15 = 0.3434, P = 0.5666) and y-axis intercept (F 1, 16 = 2.162, P = 0.1608), see Figure 4E. For the relationship between cocaine activity and cocaine activity NBA, linear regression analysis revealed that there were no differences between males and females for slope (F 1, 15 = 1.171, P = 0.2963) and y-axis intercept (F 1, 16 = 0.6221, P = 0.4418), see Figure 4F.

**Figure 4:**
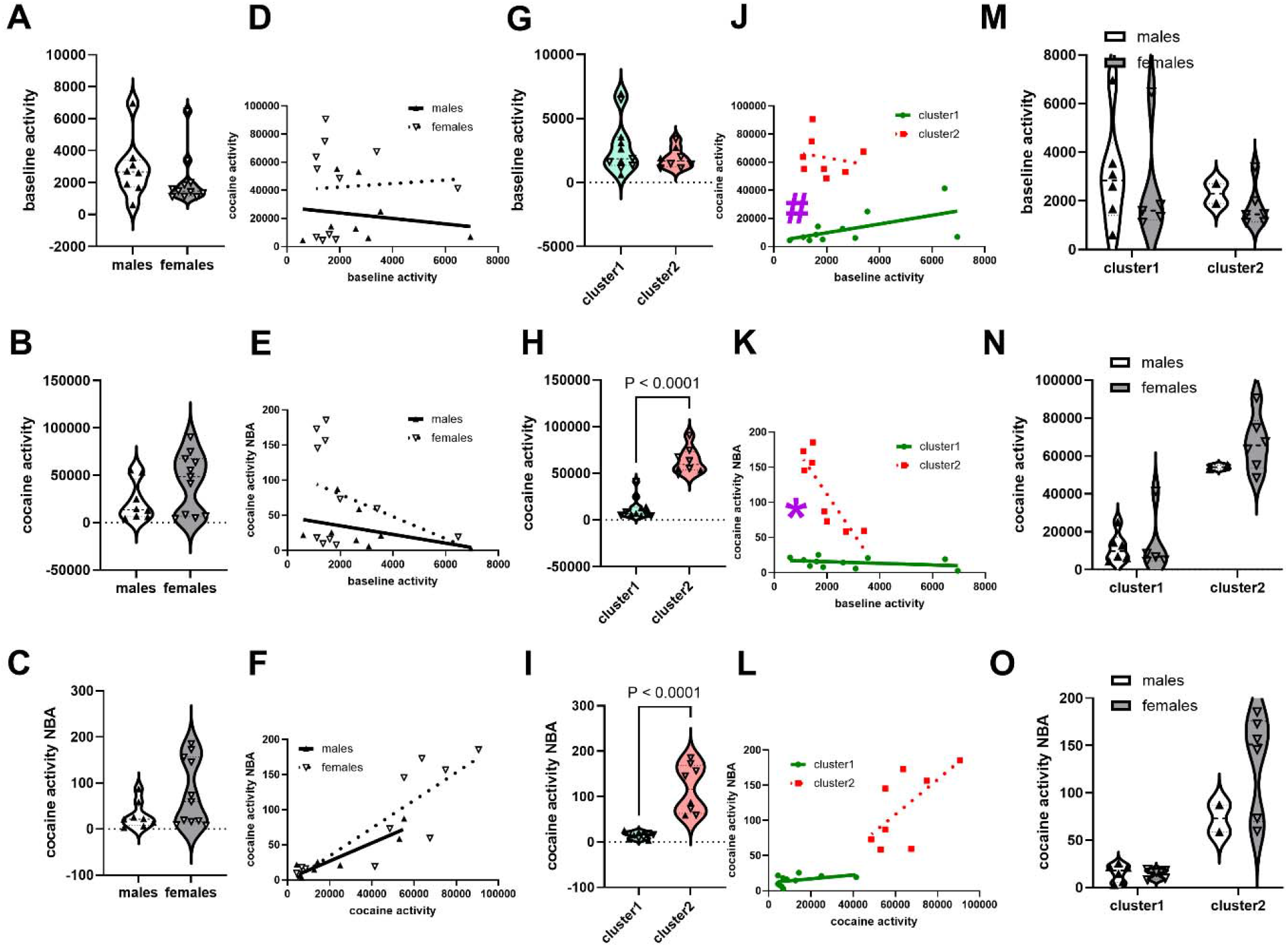
The behavioral diversity between clusters exceeds biological sex differences: cocaine group. Fig A-C show comparisons between males and females for baseline activity, cocaine activity and cocaine activity NBA, respectively, in accordance with the current model. Fig D-F, respectively, show the relationships between (baseline activity and cocaine activity), (baseline activity and cocaine activity NBA), and (cocaine activity and cocaine activity NBA), for males and females, respectively. Fig G-I show comparisons between MISSING model-identified cluster1 and cluster2 for the same variables as in Fig A-C. Fig J-L, respectively, show the relationships between baseline activity and cocaine activity, baseline activity and cocaine activity NBA, and cocaine activity and cocaine activity NBA, for cluster1 and cluster2. Fig M-O show comparisons of sex within and between clusters to determine if there was a SEX by cluster interaction (MISSING model). There were no differences between males and females for any variables/ variable relationships (Fig A-F). There were differences in variables/ variable relationships between clusters (Fig H-K), except for the baseline activity (Fig G) and for the relationship between cocaine activity and cocaine activity NBA (Fig L, not significant [but close] for slope at P = 0.06). For all the variables, Two-way ANOVA revealed no SEX by cluster interaction and no main effect of SEX (Fig M-O), but a main effect of cluster (P < 0.0001) for baseline activity (Fig M) and cocaine activity (Fig N) but not for cocaine activity NBA (Fig O). In summary, there were more significant differences for variables/ variable relationships between clusters than between biological sex. The * and # show significant differences (P < 0.05) for slope and y-axis intercepts, respectively.

#### Cluster1 versus cluster2

The baseline activity of cluster1 (n = 6 males, n = 5 females) and cluster2 (n = 2 males, n = 6 females) were 2805 ± 638 and 1893 ± 284 cm, respectively, and unpaired t-tests revealed no difference between males and females with regards to this variable (P = 0.2657, Figure 4G). The cocaine activity of cluster1 and cluster2 were 12308 ± 3443 and 63593 ± 4914 cm, respectively. Comparisons of average cocaine activity using unpaired t-tests revealed significant differences between cluster1 and cluster2 (P < 0.0001, Figure 4H). The cocaine activity NBA of cluster1 and cluster2 were 14.84 ± 2.19 and 117.39 ± 18.76, respectively, with significant differences between clusters (P < 0.0001, Figure 4I). For the relationship between baseline activity and cocaine activity, linear regression analysis revealed that there were no differences between cluster1 and cluster2 for slope (F 1, 15 = 1.157, P = 0.2992) but there were significant differences between these clusters with regards to the y-axis intercept (F 1, 16 = 84.36, P < 0.0001), see Figure 4J. For the relationship between baseline activity and cocaine activity NBA, linear regression analysis revealed significant differences between cluster1 and cluster2 for slope (F 1, 15 = 29.61, P < 0.0001, Figure 4K). For the relationship between cocaine activity and cocaine activity NBA, linear regression analysis revealed that there were no differences between clusters for slope (F 1, 15 = 4.151, P = 0.0597, almost significant) and y-axis intercept (F 1, 16 = 0.8290, P = 0.3761), see Figure 4L.

To determine if there were sex differences when we account for the clusters, we employed a Two-way ANOVA with factors SEX (males and females) and cluster (cluster1, cluster2). For baseline activity (Figure 4M), we detected no SEX × cluster interaction (F 1, 15 = 0.0006860, P = 0.9794), no main effect of cluster (F 1, 15 = 0.6750, P = 0.4242) and no main effect of SEX (F 1, 15 = 0.3992, P = 0.5370). For cocaine activity (Figure 4N), we detected no SEX × cluster interaction (F 1, 15 = 0.7245, P = 0.4080), no main effect of SEX (F 1, 15 = 1.232, P = 0.2845), but a main effect of cluster (F 1, 15 = 55.65, P < 0.0001). Likewise, for cocaine activity NBA (Figure 4O), we detected no SEX × cluster interaction (F 1, 15 = 3.533, P = 0.0797), and no main effect of SEX (F 1, 15 = 3.277, P = 0.0903), but a significant effect of cluster (F 1, 15 = 29.89, P < 0.0001).

### Model comparisons: locomotor activity time course (saline and cocaine groups)

#### Males versus Females

For locomotor activity (distance traveled-cm) for the saline and cocaine groups, we analyzed the time course data using Two-way repeated measures ANOVA with factors SEX (2 levels: males, females) and time (12 levels: time 0-120 minutes, twelve 10-minute bins including 3 baseline and 9 post-injection time points). For both groups, there was a main effect of time (P < 0.0001). For the saline group, we had n = 12 males and n = 11 females. Analysis revealed no SEX × time interaction (F 11, 242 = 1.383, P = 0.1814) and no main effect of SEX (F 1, 22 = 0.6708, P = 0.4215), see Figure 5A. Because there was no interaction, we did not proceed with post hoc tests.

**Figure 5:**
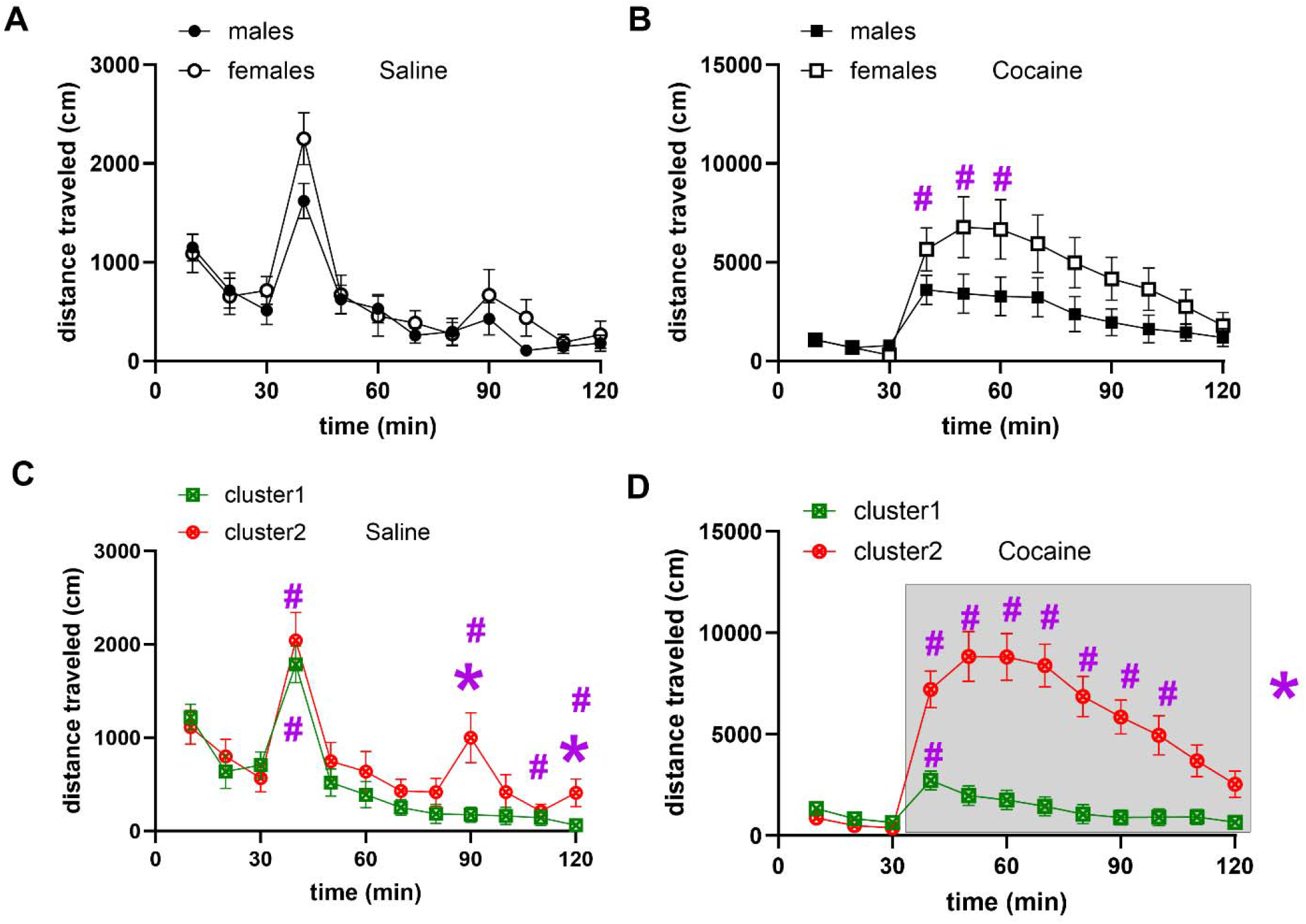
The behavioral diversity between clusters exceeds biological sex differences: time course data for saline and cocaine groups. Fig A-B represent time courses for saline and cocaine groups, respectively, where groups are designated by biological sex (current model). Fig C-D represent time courses for saline and cocaine groups, respectively, where groups are designated by behavioral clusters (MISSING model). For all analyses, there was a main effect of time (P < 0.0001). For saline group males versus females (Fig A), there was no SEX × time interaction (F 11, 242 = 1.383, P < 0.05). For cocaine group males versus females (Fig B), there was a SEX × time interaction (F 11, 198 = 2.230, P = 0.0143). For saline group cluster1 versus cluster2 (Fig C), there was a SEX × time interaction (F 11, 231 = 1.910, P = 0.0389). For cocaine group cluster1 versus cluster2 (Fig D), there was a SEX × time interaction (F 11, 209 = 19.75, P < 0.0001). The * and # show significant differences (P < 0.05) from group and from injection time point (time = 30 min), respectively, following Tukey’s post hoc tests.

For the cocaine group, we had n = 8 males and n = 11 females. Analysis revealed a SEX × time interaction (F 11, 198 = 2.230, P = 0.0143), but no main effect of SEX (F 1, 18 = 2.683, P = 0.1188). Tukey’s post hoc tests revealed no differences at any time point between biological sex groups (males versus females) but differences between pre-injection time point (30 min) versus post injection time points 40-60 min for females (Figure 5B). There were no differences between pre-injection time point (30 min) and any other time points for males.

#### Cluster1 versus cluster2

The MISSING model identified two clusters of the saline group: cluster1 had n = 12 subjects (n = 7 males and n = 5 females) and cluster2 had n = 11 subjects (n = 5 males and n = 6 females), see Figure 2A-B. For the cocaine group, MISSING model identified two clusters: cluster 1 (n = 7 males and n = 5 females) and cluster2 (n = 5 males and n = 6 females), see Figure 2C-D. We analyzed the time course data with Two-way repeated measures ANOVA with factors cluster (2 levels: cluster1, cluster2) and time (12 levels: time 0-120 minutes, twelve 10-minute bins). For both saline and cocaine groups, there was a main effect of time (P < 0.0001).

The analysis revealed a cluster × time interaction (F 11, 231 = 1.910, P = 0.0389), but no main effect of cluster (F 1, 21 = 1.999, P = 0.1720). Tukey’s post hoc tests revealed differences between clusters at time point 90 min and 120 min (Figure 5C). Tukey’s post hoc tests also revealed differences between pre-injection time point (30 min) versus post injection time points 40, 90, 110 and 120 min for cluster 1. There were differences between pre-injection time point (30 min) and only time point at 40 min for cluster2 (Figure 5C).

The analysis revealed a cluster × time interaction (F 11, 209 = 19.75, P < 0.0001) and of cluster (F 1, 19 = 32.13, P < 0.0001). Tukey’s post hoc tests revealed differences between cluster1 and cluster2 at all time points post-injection (40-120 min) (Figure 5C). There were differences between pre-injection time point (30 min) and only time point at 40 min for cluster1. Tukey’s post hoc tests also revealed differences between pre-injection time point (30 min) versus post injection time points 40-100 min for cluster2 (Figure 5C).

### Assessments of sex differences within and between clusters: time course analysis

We employed Two-way repeated measures ANOVA with factors SEX and time. For all analysis there was a main effect of time (P < 0.05). For the saline group, there were no SEX × time interactions for comparisons between males and females within cluster1 (F 11, 110 = 0.7179, P = 0.7192, Figure 6A) and cluster2 (F 11, 99 = 1.035, P = 0.4224, Figure 6B). There were no SEX × time interactions for comparisons between males in cluster1 and females in cluster2 (F 11, 121 = 1.484, P = 0.1458, Figure 6C) and between males in cluster2 versus females in cluster1 (F 11, 88 = 1.860, P = 0.0558, Figure 6D). For the cocaine group, there were no SEX × time interactions for comparisons between males and females within cluster1 (F 11, 99 = 0.7151, P = 0.7216, Figure 6E) and cluster2 (F 11, 66 = 0.4546, P = 0.9241, Figure 6F). However, there were SEX × time interactions for comparisons between males in cluster1 versus females in cluster2 (F 11, 110 = 15.32, P < 0.0001, Figure 6G) and between males in cluster2 versus females in cluster1 (F 11, 55 = 16.22, P < 0.0001, Figure 6H).

**Figure 6:**
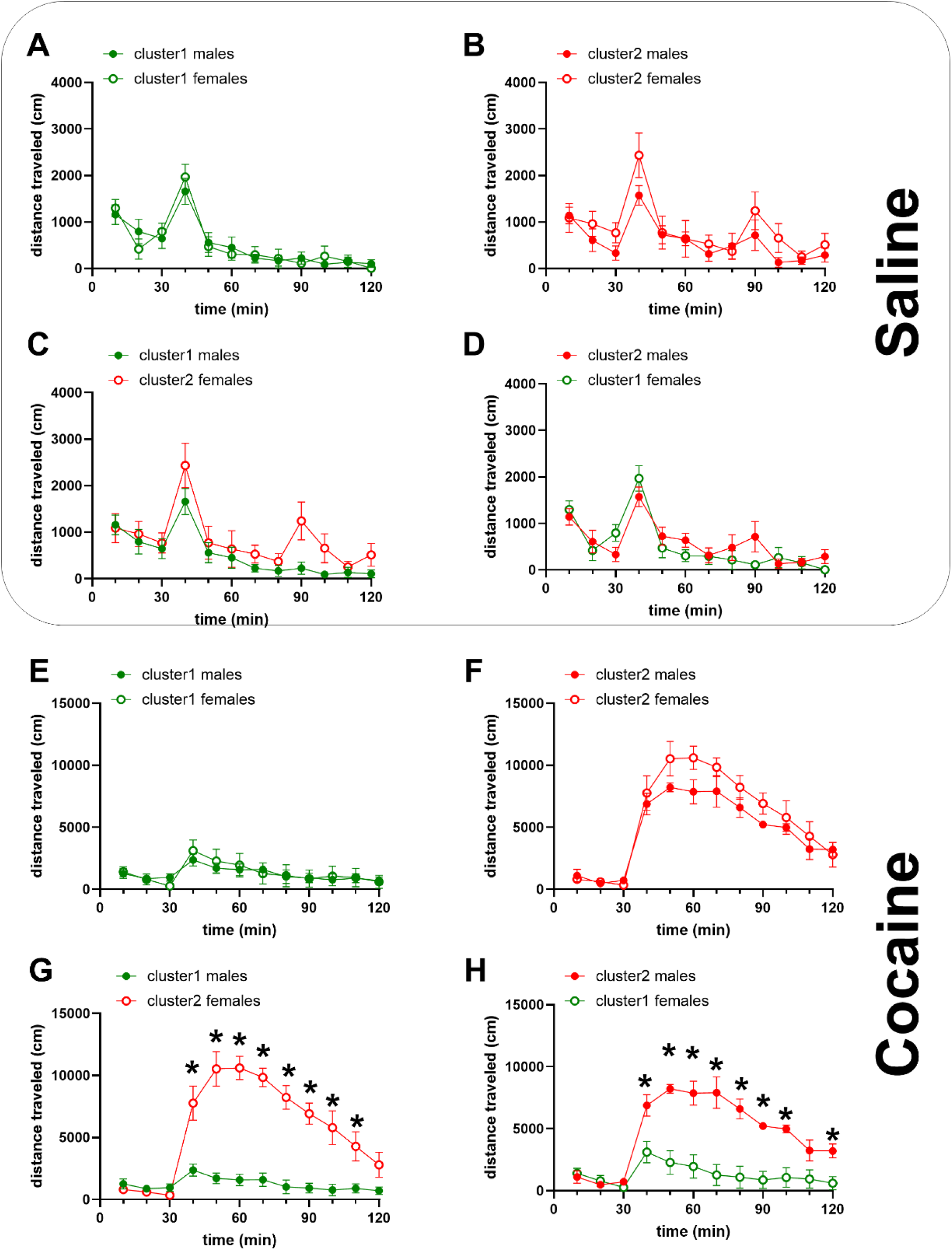
Differences between males and females, when observed, are mostly dependent on behavioral cluster differences not biological sex: time course data for saline and cocaine groups. Fig A-D represent locomotor activity time courses for the saline group while Fig E-H represent time courses for the cocaine group. For all comparisons, Two-way repeated measures ANOVA revealed a main effect of time (P < 0.05). There was no SEX × time interaction (p > 0.05) for comparisons between males and females within the same clusters for saline or cocaine groups (Fig A-B, E-F). For the saline group, there was no SEX × time interaction for cluster1 males versus cluster2 females (F 11, 121 = 1.484, P = 0.1458, Fig C) and cluster2 males versus cluster1 females (F 11, 88 = 1.860, P = 0.0558, Fig D). However, for the cocaine group, there was a significant SEX × time interaction for cluster1 males versus cluster2 females (F 11, 110 = 15.32, P < 0.0001, Fig G) and cluster2 males versus cluster1 females (F 11, 55 = 16.22, P < 0.0001, Fig H). The * show significant differences (P < 0.05) between group when SEX × time interaction was followed up with Tukey’s post hoc tests. In summary, sex differences are most significant when males and females from different, not same, clusters are compared.

### Summary of Results

We compared the current model with the MISSING model (Figure 1), specifically to determine which grouping strategy (males versus females for the current model, cluster versus cluster for the MISSING model) reveals more distinct groups. Normal mixtures clustering yielded 2 clusters for saline and cocaine group (Figure 2). For the saline group (Figure 3) and the cocaine group (Figure 4), cluster1 was more different from cluster2 than males were different from females. For locomotor activity time course data (Figure 5) for saline and cocaine injected subjects, cluster1 was more different from cluster2 than males were different from females. There were no sex differences when males and females within the same cluster were compared (Figure 6A-B, E-F). When sex differences were observed, it was because males and females from different clusters were compared (Figure 6G-H), suggesting that even the observed significant differences between males and females are due to a behavioral group-related effect and not due to biological sex *per se*.

## Discussion

Our hypothesis was that group identity (revealed by the MISSING model) would be a more effective grouping strategy than biological sex (current model). To test this hypothesis, we compared these models and determined group identity (consisting of both males and females), was more effective than biological sex (males versus females) in identifying distinct behavioral groups. Our results confirm our hypothesis and further validate the MISSING model, see (Job, 2024; Showell and Job, 2024; Tigano and Job, 2024).

As with previous studies (Job, 2024; Showell and Job, 2024; Tigano and Job, 2024), the MISSING model confirmed that 1) there were no sex differences when males and females within a cluster were compared, and 2) when sex differences were detected, it was because of comparisons of males and females from different clusters. By showing that sex differences are most pronounced when we compare males and females with different behavioral group identities, detecting sex differences using the current model may actually be detecting group identity differences unrelated to biological sex per se. As such, characterizing the biochemical mechanisms governing these differences and ascribing them to biological sex may not be the most effective way to understand the mechanisms governing sex differences. *We propose that to understand sex differences in behavior, it is essential to compare males and females that belong to the same behavioral cluster instead of comparing males and females generally as is done in the current model*. This new approach would compare males and females with similar behavior (similar baseline) and, as such, any differences in molecular mechanisms despite these behavioral similarities are more likely to be more accurate representations of the mechanism governing sex differences, if any, for such behavior.

We detected two clusters consisting of males and females for the saline group. This suggests that the MISSING model is sensitive enough to identify clusters with differential behavioral responses to injections of vehicle controls. These groups may represent a diversity in individual responses to the effects of stressors on behavior. If control groups can display this behavioral diversity, the impact of this observation must be considered if we are to understand drug behavior relative to controls (this is not currently done).

Limitations of the study include the number of subjects used, particularly for the male cocaine group. However, despite this limitation, when we combine the results gleaned from this study with the results from previous studies see (Job, 2024; Showell and Job, 2024; Tigano and Job, 2024), we achieve the goals of further validating the MISSING model. Another limitation is that the estrous phase of the females was not assessed. However, a previous report revealed that estrus phase does not appear to be predict behavioral clusters (Tigano and Job, 2024). This study does, however, reveal the limitations of utilizing the current model to understand sex differences by revealing that while attempting to distinguish groups based on biological sex, one can simultaneously be blind to valuable information regarding behavioral diversity which may be potentially more relevant to behavioral data than biological sex.

In summary, the MISSING model is more effective than the current model that employs sex as a biological variable in identifying behavioral diversity. The MISSING model also proposes a new approach for understanding sex differences by comparing males and females within the same behavioral cluster instead of grouping by biological sex generally. This represents a paradigm shift from the status quo and has the potential to truly advance the field of sex differences research.

## Acknowledgements

The authors wish to acknowledge Michael J Kuhar in whose laboratory MOJ conducted the behavioral experiments. All authors contributed to data analysis and to the writing of the manuscript. MOJ designed and conducted behavioral experiments and statistical analysis. The authors acknowledge the support of NIH grants DA15162 and DA015040. Also this work was funded by the National Center of Research Resources P51RR165 and was supported by the Office of Research Infrastructure Programs/OD P51OD11132. This work was also supported by the Francis Lax Fund for Faculty Development at Rowan University. This work was also supported by startup funds from Rowan University, Camden, New Jersey.

